# Bacteriophage diversity declines with COPD severity in the respiratory microbiome

**DOI:** 10.1101/2025.06.16.659870

**Authors:** Ryan A. Cook, Alise J. Ponsero, Andrea Telatin, Yuqiong Yang, Zhenyu Liang, Fengyan Wang, Rongchang Chen, Zhang Wang, Evelien M. Adriaenssens, Martha RJ. Clokie, Andrew D. Millard, Christopher E. Brightling

## Abstract

Chronic obstructive pulmonary disease (COPD) severity correlates with airway microbial dysbiosis, yet bacteriophage roles remain unexplored. We characterised the lung virome by re-analysing 135 sputum metagenomes from 99 COPD patients and 36 healthy controls. We identified 1,308 viral operational taxonomic units, revealing progressively lower viral diversity correlating with disease severity. Whilst viral and bacterial diversity typically showed strong positive correlations, patients with frequent exacerbations uniquely exhibited decoupled viral-bacterial relationships, indicating disrupted ecological dynamics. Phages infecting anaerobic oral bacteria showed disproportionately lower abundance—*Porphyromonas* phages were 40-fold less abundant despite only 4-fold lower bacterial abundance—whilst pathogen-associated phages showed no significant differences. We detected virulence factor-encoding phages, including two *neuA*-carrying *Haemophilus* phages in 7.4% of *Haemophilus*-colonised patients, associated with 82-fold higher bacterial abundance. These findings establish altered bacteriophage ecology as an unrecognised feature of COPD pathobiology, with differential phage-bacteria relationships that reshape lung microbial ecosystems, offering new perspectives for microbiome-targeted interventions.

## Background

Chronic obstructive pulmonary disease (COPD) ranks among the leading causes of mortality worldwide, responsible for over three million deaths annually^1^. This heterogeneous disease encompasses emphysema and chronic bronchitis, with patients experiencing varied clinical trajectories. Disease severity is assessed through the Global Initiative for Chronic Obstructive Lung Disease (GOLD) staging system, grading airflow limitation from mild (GOLD 1) to very severe (GOLD 4)^2^. However, recognising that airflow limitation alone inadequately predicts clinical outcomes, the ABE assessment tool stratifies patients by symptom burden and exacerbation history—Group A (low symptoms/low exacerbations), Group B (high symptoms/low exacerbations), and Group E (frequent exacerbations)^2^. This emphasis on exacerbation frequency reflects its critical role as an independent predictor of mortality and disease progression. Beyond clinical classifications, COPD manifests distinct inflammatory endotypes influencing treatment responses. Neutrophilic inflammation (elevated sputum neutrophils) associates with bacterial colonisation and poor corticosteroid response, whilst eosinophilic inflammation (elevated sputum eosinophils) predicts better corticosteroid outcomes^3^.

The bacterial component of the lung microbiome plays a well-established role in COPD pathogenesis, with dysbiosis linked to disease progression, exacerbations, and therapeutic responses^4-6^. However, the viral component remains unexplored beyond eukaryotic respiratory viruses in acute exacerbations^7^. This oversight is striking given bacteriophages’ profound influence on microbial communities across all ecosystems^8-10^.

Bacteriophages shape bacterial communities through diverse mechanisms beyond predation. Lytic phages kill hosts without integration into the host genome, regulating populations through "Kill-the-Winner" dynamics^11^. Temperate phages integrate as prophages into the host genome, often carrying virulence factors that can transform benign bacteria into pathogens^12,13^. Environmental stresses—including antibiotics, inflammation, or nutrient limitation—can trigger prophage induction, reshaping microbial communities^14,15^.

In cystic fibrosis, which shares pathophysiological features with COPD, bacteriophages directly impact clinical outcomes. Filamentous phages impair mucociliary clearance, promote antibiotic resistance, and correlate with worse prognosis^16,17^, suggesting phages may similarly influence COPD pathogenesis.

The recent publication of comprehensive COPD metagenomes by Yan et al.^18^ revealed airway microbiome alterations associated with neutrophilic inflammation and identified microbial metabolites linked to disease severity. However, the previous analysis focused on bacteria, leaving viral components unexamined. Here, we applied state-of-the-art bioinformatic approaches to study the DNA virome of these data, revealing alterations in viral ecology associated with COPD. We demonstrate progressive viral diversity loss correlating with disease severity, identify decoupled viral-bacterial relationships in frequent exacerbators (Group E), and detect virulence-encoding phages associated with dramatically elevated pathogen abundance. These findings establish the virome as a previously unrecognised component of COPD pathobiology with implications for understanding disease mechanisms and developing novel therapeutics.

## Results

### Study Cohort and Viral Catalogue Overview

We re-analysed 135 sputum metagenomes from 99 COPD patients and 36 healthy controls originally published by Yan et al. The cohort spanned the clinical spectrum of COPD: GOLD stages 1-4 (n=21, 40, 29, and 9 respectively, Figure 1A), ABE groups A (n=48, low symptoms/low exacerbations), B (n=29, high symptoms/low exacerbations), and E (n=20, high exacerbations) (Figure 1B), and inflammatory endotypes including neutrophilic (n=31), eosinophilic (n=11), mixed granulocytic (n=50), and paucigranulocytic (n=5) (Figure 1C). Two patients lacked metadata for classification.

**Figure 1.**
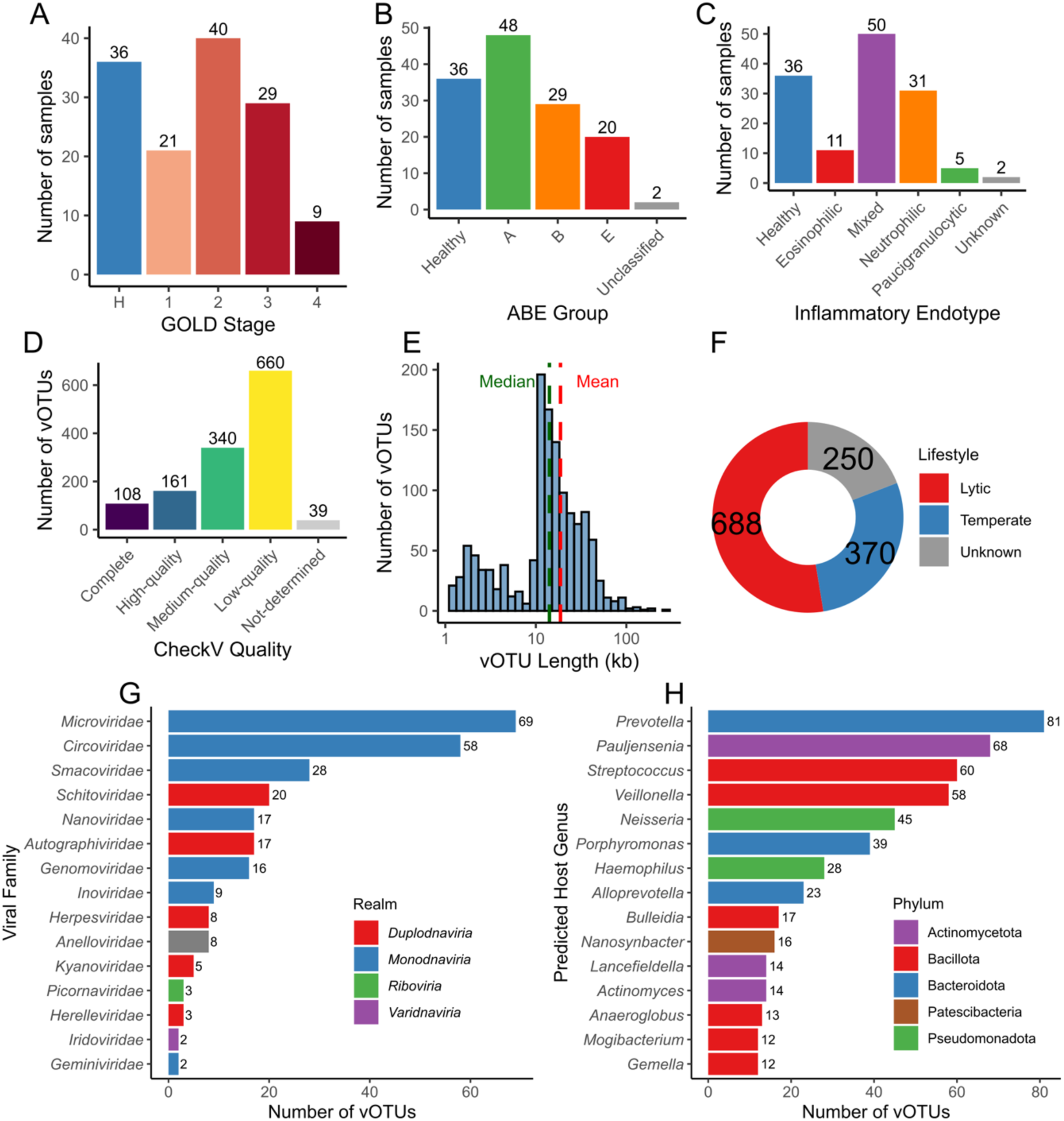
Overview of study cohort and viral catalogue characteristics. (**A**) Distribution of samples across GOLD stages, including healthy controls (H) and GOLD stages 1-4. (**B**) Distribution of samples across ABE classification groups: Healthy, Group A (low symptoms/low exacerbations), Group B (high symptoms/low exacerbations), Group E (frequent exacerbations), and Unclassified. (**C**) Distribution of samples across inflammatory endotypes. (**D**) CheckV quality assessment of 1,308 viral operational taxonomic units (vOTUs) categorised as Complete, High-quality, Medium-quality, Low-quality, or Not-determined. (**E**) Length distribution of vOTUs displayed on a log10 scale with mean (red dashed line) and median (green dashed line) indicated. (**F**) Top 15 most frequently predicted viral families, coloured by realm classification. (**G**) Top 15 most frequently predicted bacterial host genera, coloured by phylum. (**H**) Predicted viral lifestyle, showing temperate, lytic, and unknown categories. Numbers above bars indicate number of samples or vOTUs for each category.

Viral mining yielded 1,308 viral operational taxonomic units (vOTUs; approximate to species, Figure 1D), ranging from 1.1-252.2 kb (median 14.1 kb, Figure 1E). Quality assessment classified 108 vOTUs as complete, 161 as high-quality (>90% predicted complete), and 340 as medium-quality (>50% predicted complete) (Figure 1G). The majority (79.3%) represented novel viral diversity, with only 271 vOTUs assignable to known families, though 892 (68.2%) could be classified as *Caudoviricetes* (tailed phages). Lifestyle predictions indicated 688 obligately lytic and 370 temperate viruses (Figure 1F), whilst host predictions linked 675 vOTUs to 88 bacterial genera, most commonly *Prevotella* (n=81), *Pauljensenia* (n=68), and *Streptococcus* (n=60) (Figure 1H).

### Progressive Viral Diversity Loss Correlates with COPD Severity

Viral alpha diversity declined significantly with disease severity across multiple clinical classifications. Shannon diversity showed progressive reduction from healthy controls (1.93 ± 1.27) through GOLD stages 1-2 (1.27 ± 1.08 and 1.33 ± 1.16) to stages 3-4 (0.83 ± 0.97 and 0.69 ± 0.89; Kruskal-Wallis χ² = 16.99, p = 0.002). Post-hoc analyses confirmed significant reductions in GOLD 3 (p = 0.001) and GOLD 4 (p = 0.010) compared to controls, suggesting a threshold effect where viral community disruption becomes pronounced in severe disease (Figure 2A). The substantial within-group variation was notable, with some patients harbouring no detectable vOTUs in their metagenomes despite adequate sequencing depth, highlighting the heterogeneous nature of viral depletion in COPD.

**Figure 2.**
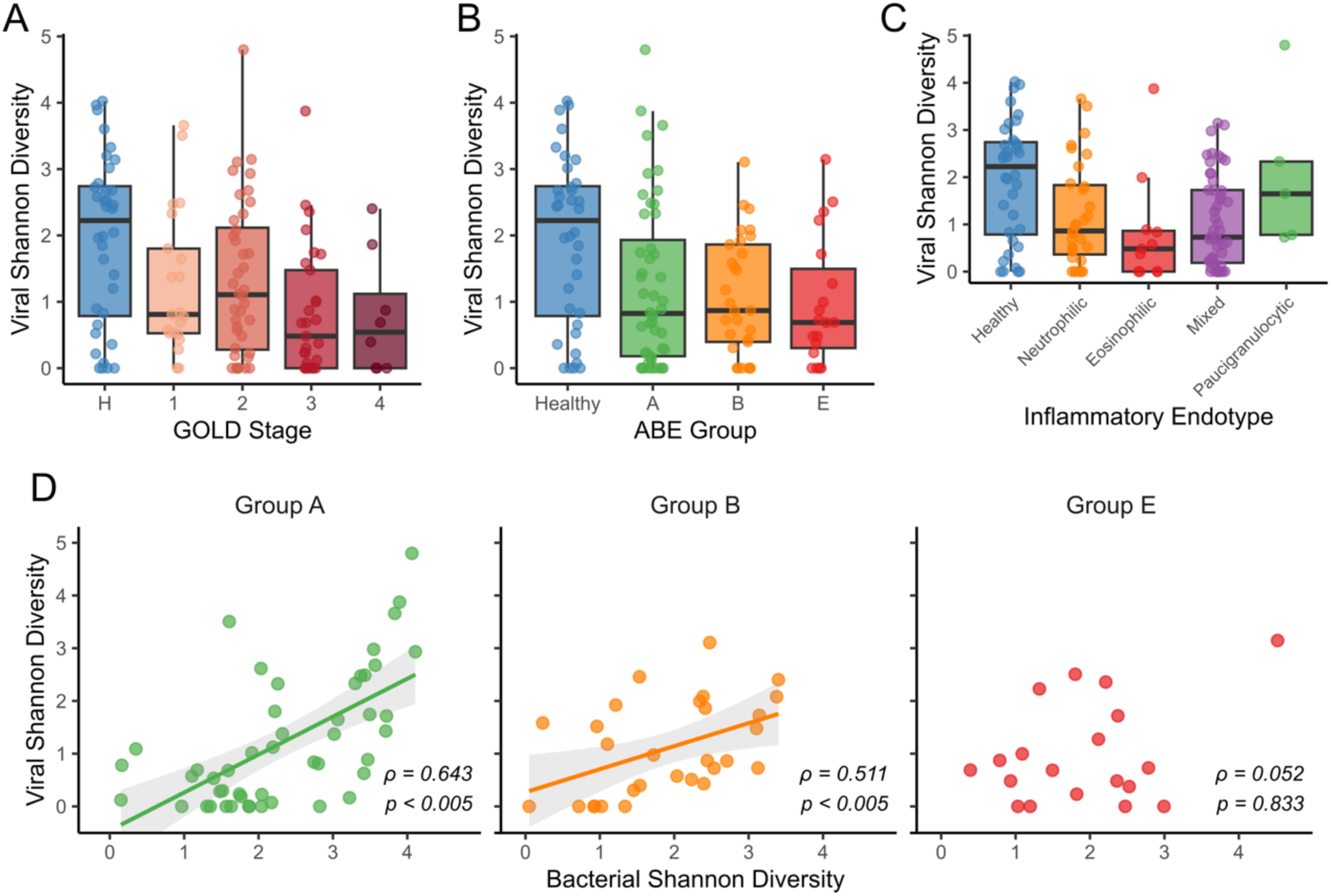
Progressive viral diversity loss and ecological decoupling in COPD. (**A**) Viral Shannon diversity across GOLD stages showing progressive decline with disease severity (n = 135; Kruskal-Wallis χ² = 16.99, df = 4, P = 0.002). (**B**) Viral Shannon diversity across ABE groups demonstrating reduced diversity in all COPD phenotypes compared to healthy controls (n = 135; Kruskal-Wallis χ² = 13.47, df = 4, P = 0.009). (**C**) Viral Shannon diversity by inflammatory endotype revealed significantly reduced diversity in eosinophilic (P = 0.017) and mixed granulocytic (P = 0.011) phenotypes compared to healthy controls (n = 130 with endotype data; Kruskal-Wallis χ² = 15.45, df = 5, P = 0.009). (**D**) Correlation between bacterial and viral Shannon diversity stratified by ABE group, demonstrating preserved ecological coupling in Groups A (ρ = 0.643, P < 0.005) and B (ρ = 0.511, P < 0.005) but complete decoupling in frequent exacerbators (Group E; ρ = 0.052, P = 0.833). For all box plots: centre line, median; box limits, upper and lower quartiles; whiskers, 1.5× interquartile range; points, individual samples. Spearman correlation coefficients (ρ) and P values shown for each group. Shaded areas represent 95% confidence intervals for linear regression lines.

All ABE groups showed reduced viral diversity compared to controls (Kruskal-Wallis χ² = 13.47, p = 0.009): Group A (1.23 ± 1.24, p = 0.019), Group B (1.10 ± 0.89, p = 0.018), and Group E (0.94 ± 0.96, p = 0.022). However, no differences existed between ABE groups (all pairwise p > 0.05), indicating viral depletion occurs independently of symptom burden or exacerbation frequency (Figure 2B). Community evenness (Inverse Simpson diversity) showed a trend toward lower values in Group E (p = 0.051), suggesting their viral communities may be dominated by fewer abundant species.

Inflammatory endotypes also showed significant differences in viral diversity (Kruskal-Wallis χ² = 15.45, p = 0.009). Patients with eosinophilic inflammation showed the most pronounced reduction in viral diversity (0.82 ± 1.17 vs 1.93 ± 1.27 in controls, p = 0.017, Figure 2C). Viral diversity negatively correlated with sputum eosinophil counts (Spearman’s ρ = -0.257, p = 0.003), as did bacterial diversity (ρ = -0.232, p = 0.007). However, mediation analysis revealed these relationships are interdependent, with 61% of the eosinophil-viral association mediated through bacterial diversity (p = 0.028), suggesting type 2 inflammation may drive coordinated suppression of both microbial domains.

Analysis of predicted viral lifestyle revealed additional endotype-specific patterns. Whilst temperate virus relative abundance remained constant across all disease classifications, obligately lytic virus proportions differed significantly by inflammatory endotype (Kruskal-Wallis χ² = 13.70, p = 0.018). Patients with a paucigranulocytic endotype showed dramatically elevated obligately lytic virus abundance (72.4% ± 17.4%) compared to all other groups: healthy controls (30.2% ± 24.2%, p = 0.004), eosinophilic (16.6% ± 15.8%, p = 0.016), mixed granulocytic (25.4% ± 25.1%, p = 0.006), and neutrophilic inflammation (24.7% ± 28.6%, p = 0.004). This 3-fold increase in obligately lytic viruses correlated strongly with overall viral community metrics (Shannon diversity: ρ = 0.671, p < 0.001; total viral load: ρ = 0.436, p < 0.001), suggesting that lytic viral activity may drive viral community richness in low-inflammation environments.

Viral load (percentage of total reads mapping to viral contigs) correlated positively with all viral diversity metrics (Shannon: ρ = 0.45, p < 0.001; richness: ρ = 0.52, p < 0.001; Inverse Simpson: ρ = 0.31, p < 0.001). Patients with high viral loads (>75th percentile, >2.1%) maintained significantly greater diversity than those with low loads (<25th percentile, <0.6%) across all metrics (Wilcoxon test, all p < 0.001), with high-load patients averaging 10-fold higher richness (41.1 ± 57.7 vs 4.1 ± 4.8 vOTUs).

### Disrupted Viral-Bacterial Ecological Relationships in Frequent Exacerbators

Viral and bacterial diversity typically showed strong positive correlations (richness: ρ = 0.704, p < 0.0001; Shannon: ρ = 0.502, p < 0.0001), with both communities declining in parallel with disease severity (Figure 2D). Healthy controls maintained the highest bacterial diversity (Shannon: 2.98 ± 0.81, 86.5 ± 63.8 genera), with all COPD groups showing significant reductions (all p < 0.001).

However, this coordinated relationship uniquely broke down in patients with frequent exacerbations (Group E), for which the correlation between viral and bacterial diversity disappeared (ρ = 0.052, p = 0.833), contrasting sharply with maintained correlations in Groups A (ρ = 0.643, p < 0.005) and B (ρ = 0.511, p < 0.005). Partial correlation analysis controlling for GOLD stage and inflammatory endotype confirmed this pattern: Groups A and B retained significant correlations (partial ρ = 0.676 and 0.442, respectively; both p < 0.05), whilst Group E remained uncorrelated (partial ρ = -0.198, p = 0.447). This decoupling in frequent exacerbators represents a distinct ecological disruption independent of disease severity or inflammatory status.

The decoupling was specific to the ABE classification. When stratified by GOLD stage, all groups except GOLD 4 (likely due to small sample size, n=8) maintained significant viral-bacterial correlations (GOLD 1: ρ = 0.542, p = 0.011; GOLD 2: ρ = 0.378, p = 0.016; GOLD 3: ρ = 0.452, p = 0.014). Similarly, all inflammatory endotypes preserved the relationship: neutrophilic (ρ = 0.507, p = 0.0036), mixed granulocytic (ρ = 0.443, p = 0.0014), and eosinophilic (ρ = 0.442, though p = 0.174 due to small sample size, n=11).

### Limited Viral Community Structuring Despite Diversity Loss

Beta diversity analysis using Bray-Curtis dissimilarity revealed subtle but significant viral community structuring by clinical phenotypes. ABE classification explained 4.5% of community variation (PERMANOVA R² = 0.045, p = 0.02, Figure 3A), with Group B (high symptoms, low exacerbations) harbouring the most distinct viral communities, differing significantly from healthy controls (R² = 0.051, adjusted p = 0.020) and Group A (R² = 0.032, adjusted p = 0.023). Surprisingly, neither Group A nor E differed from controls (p = 0.610 and 0.706, respectively), and Groups A and E were virtually indistinguishable (R² = 0.008, p = 0.974).

**Figure 3.**
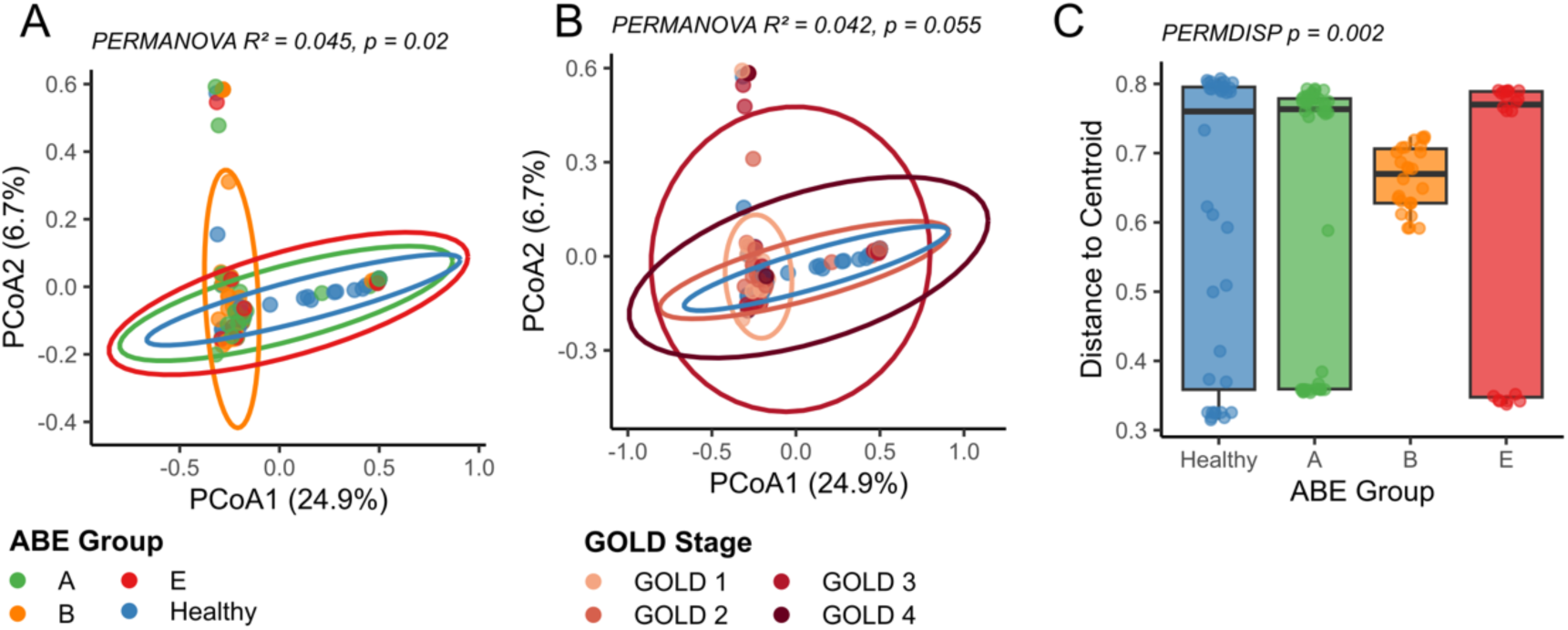
Limited viral community structuring across COPD clinical phenotypes. (**A**) Principal coordinates analysis (PCoA) of viral community composition based on Bray-Curtis dissimilarity, coloured by ABE classification. Ellipses represent 95% confidence intervals. (**B**) PCoA of viral communities coloured by GOLD stage. (**C**) Beta-dispersion analysis showing distances to group centroids for ABE classifications. Box plots show median and interquartile range; individual points represent samples. PERMANOVA and PERMDISP statistics are shown on respective panels. n = 36 healthy controls, n = 36 Group A, n = 28 Group B, n = 29 Group E for ABE classification; n = 36 healthy, n = 28 GOLD 1, n = 55 GOLD 2, n = 14 GOLD 3, n = 8 GOLD 4 for GOLD stages.

Overall COPD status did not create a distinct viral signature at the vOTU level (R² = 0.011, p = 0.132), though clinical subgroups within COPD showed significant differentiation (R² = 0.047, p = 0.036). Significant heterogeneity in multivariate dispersion (PERMDISP: ABE groups F = 5.19, p = 0.002; endotypes F = 3.77, p = 0.003, Figure 3C) indicated some groups harbour more variable viral communities, potentially reflecting different stages of ecological stability. Neither GOLD classification (R² = 0.042, p = 0.055, Figure 3B) nor inflammatory endotypes (R² = 0.050, p = 0.056) showed significant effects.

In contrast, bacterial communities showed stronger clinical structuring: ABE groups (R² = 0.067, p = 0.005), GOLD stages (R² = 0.066, p = 0.004), inflammatory endotypes (R² = 0.082, p = 0.003), and COPD vs healthy status (R² = 0.041, p = 0.003) all showed significant associations. Both communities experienced diversity loss, but only bacterial communities maintained clear associations with clinical phenotypes.

Although viral communities showed weak clinical structuring, they correlated significantly with bacterial community composition (Mantel r = 0.271, p = 0.001). This relationship persisted after controlling for disease status (r = 0.276, p = 0.001) or GOLD stage (r = 0.270, p = 0.001). Procrustes analysis confirmed significant congruence between viral and bacterial ordinations (correlation = 0.383, p = 0.001), with communities sharing approximately 15% of compositional variation despite their divergent clinical associations (correlation² = 0.147). Environmental fitting identified patient-reported symptom burden (CAT score) as the only clinical variable significantly associated with viral structure (r² = 0.108, p = 0.003), whilst objective measures including FEV1, BMI, and inflammatory markers showed no significant associations.

### Selective Depletion of Commensal-Associated Phages

Analysis of host-specific viral abundances revealed unexpected patterns. The key COPD pathogenic genera—*Streptococcus* (4.4% mean relative abundance, 68.1% prevalence), *Haemophilus* (3.4%, 50.4%), and *Moraxella* (2.0%, 11.1%, significantly elevated in COPD, p = 0.014)—were predicted as hosts for 90 vOTUs total. However, viruses infecting these classic COPD associated pathogens showed no significant differential abundance between groups after multiple testing correction.

Instead, consistent depletion occurred for viruses infecting oral and anaerobic genera (Figure 4A). *Porphyromonas* viruses showed the strongest association (Kruskal-Wallis χ² = 25.4-26.9, p < 0.001 across all comparisons), with 40-fold reduction in COPD (mean 25,792 vs 636 CPM) and prevalence dropping from 36.1% to 4.0%. Crucially, this viral depletion exceeded bacterial depletion—*Porphyromonas* bacterial abundance decreased only 4-fold (8.49% to 1.93%), suggesting virus-specific loss beyond host availability (Figure 4B).

**Figure 4.**
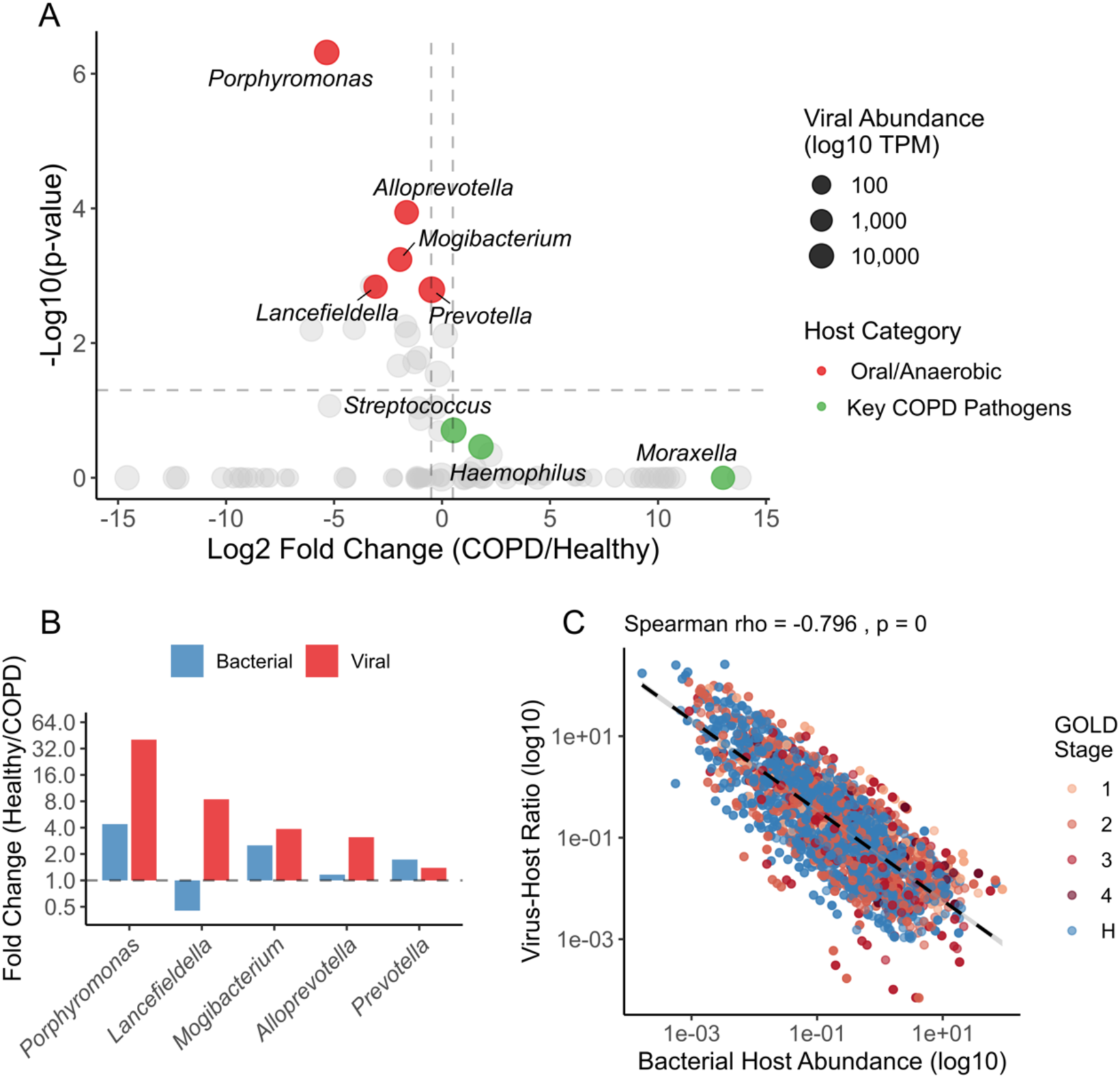
Selective depletion of oral and anaerobic bacteria-associated phages in COPD. (**A**) Volcano plot showing differential abundance of phages, grouped by predicted bacterial host between COPD patients (n=99) and healthy controls (n=36). Points represent individual bacterial genera, with size indicating mean viral abundance (log10 CPM) and colour indicating host category. Dashed lines indicate log2 fold change = ±0.5 and adjusted p = 0.05 thresholds. (**B**) Comparison of viral and bacterial depletion for oral and anaerobic genera. Bars show fold change (healthy/COPD) for viral abundance and corresponding bacterial host abundance. (**C**) Virus-host ratio (VHR) plotted against bacterial host abundance across all samples (n=111 with sufficient data), coloured by GOLD stage. Dashed line shows linear regression fit. Spearman correlation coefficient and p-value are indicated. Both axes are log10-scaled.

Similar disproportionate patterns occurred for *Alloprevotella* viruses (3-fold reduction despite only 1.2-fold bacterial change), *Mogibacterium* viruses (3.9-fold reduction vs 2.5-fold bacterial depletion), and *Lancefieldella* viruses. Of the differentially abundant host-grouped viruses, only *Prevotella* showed proportional changes (1.4-fold viral reduction, 1.7-fold bacterial reduction). These depletion patterns followed disease severity gradients, with progressive loss across GOLD stages and ABE classifications, suggesting selective disruption of virus-bacteria relationships for commensal organisms.

Virus-host ratio (VHR) analysis, calculated by dividing viral abundance by corresponding bacterial host abundance, encompassed 3,619 observations across 45 host genera. The strong negative correlation between VHR and bacterial abundance (overall ρ = -0.796, p < 0.001, Figure 4C) aligns with "Kill-the-Winner" ecological dynamics, where viral predation preferentially targets abundant bacterial populations. This pattern persisted across all clinical classifications: ABE groups (ρ = -0.816 to -0.663, all p < 0.001), GOLD stages including controls (ρ = -0.591 to -0.853, all p < 0.01), and inflammatory endotypes (ρ = -0.701 to -0.906, all p < 0.05). The strongest correlations occurred in patients with eosinophilic inflammation (ρ = -0.892) and early disease (GOLD 1: ρ = -0.853), whilst the weakest appeared in severe disease (GOLD 4: ρ = -0.591), suggesting altered ecological dynamics in late-stage disease.

### Acute Viral Events: Eukaryotic Infections and Bacteriophage Blooms

Five patients exhibited extreme viral loads (>10% metagenomic reads mapping to vOTUs), revealing distinct acute phenomena beyond the chronic viral depletion patterns (Figure 5A-B). Two harboured active eukaryotic viral infections: Human mastadenovirus B (patient K171, 47.8% viral load, 78.1% relative abundance, Figure 5E), a known respiratory pathogen, and a novel picorna-like virus most similar to a swan-associated virus (patient K170, 16.1% viral load, 91.0% relative abundance, figure 5F). As picornaviruses are RNA viruses, detection in a DNA metagenome likely resulted from reverse transcription activity during amplification prior to sequence library preparation.

**Figure 5.**
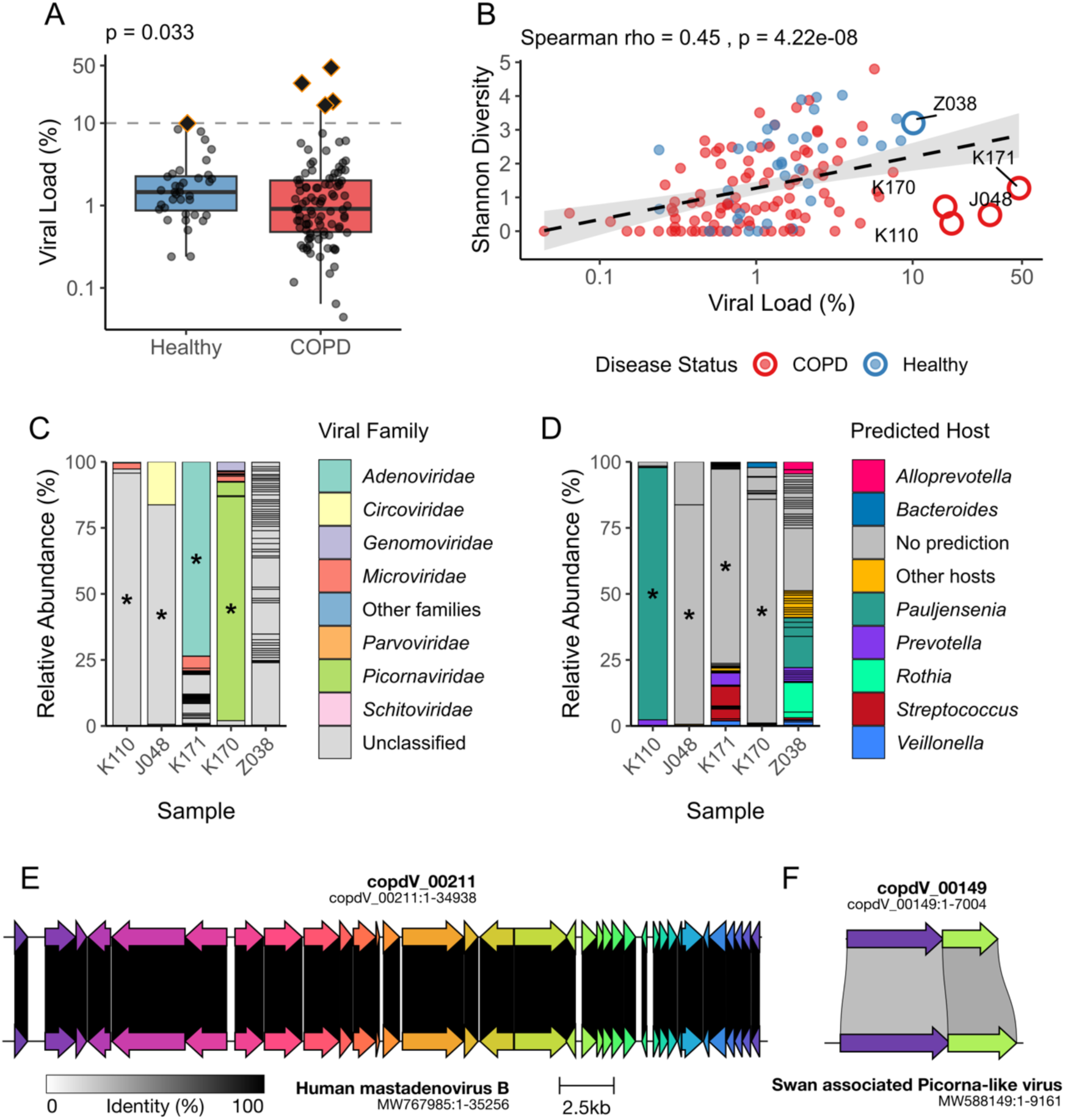
Acute viral events drive extreme viral loads in the respiratory microbiome. (**A**) Viral load comparison between healthy controls and COPD patients (y axis in log scale). Box plots show median and interquartile range, with individual samples shown as points. Diamonds indicate extreme viral load cases (>10%, dashed line). Statistical significance determined by Wilcoxon rank-sum test. (**B**) Relationship between viral load and Shannon diversity. Points are coloured by disease status, with extreme cases (>10%) highlighted with coloured borders and labelled. Dashed line shows linear regression with 95% confidence interval. Spearman correlation coefficient and p-value shown. (**C**,**D**) Composition of extreme viral load samples. Stacked bar charts show relative abundance of vOTUs in five samples with >10% viral load, coloured by (**C**) viral family or (**D**) predicted bacterial host genus. Each segment represents an individual vOTU with black borders separating vOTUs within the same colour category, with the asterisk indicating a dominant vOTU. (**E**,**F**) Synteny comparisons of dominant vOTUs from suspected viral infections. Clinker alignments show genomic organisation and similarity between (**E**) vOTU from patient K171 and Human mastadenovirus B (GenBank: MW767985), and (**F**) vOTU from patient K170 and a swan-associated picorna-like virus (GenBank: MW588149). ORFs are colour-coded by gene cluster, with grey lines indicating pairwise alignments between homologous sequences. Line intensity reflects percentage identity of aligned regions.

Two others showed "phage blooms"—dominance by single bacteriophages (Figure 5C-D). Patient K110 (17.7% viral load) was dominated by a ∼52 kb lytic phage (95.5% relative abundance) with CRISPR spacer matches to *Pauljensenia*, whilst patient J048 (31.2% viral load) contained a dominant ∼80 kb temperate phage of unknown host (83.1% relative abundance). Both belonged to class *Caudoviricetes* but lacked family-level classification. These blooms resulted in extremely low diversity (five and three vOTUs respectively, Shannon = 0.23 and 0.48), suggesting periodic viral population collapses and expansions that may reflect prophage induction events or lytic phage amplification during bacterial blooms.

### Virulence-Associated Phages Show Strong Host Coupling

Screening against the Virulence Factor Database identified four vOTUs carrying bacterial virulence genes, targeting *Haemophilus* (n=2), *Streptococcus* (n=1), and *Escherichia* (n=1). The virulence factors included *neuA* (N-acylneuraminate cytidylyltransferase, involved in lipooligosaccharide biosynthesis and immune evasion)^19^, *lytA* (autolysin, contributing to pneumococcal pathogenesis)^20^, and genes for lipopolysaccharide modification (*gtrA*, *gtrB*) that can alter bacterial surface antigens^21,22^. All four vOTUs were predicted to be temperate and may therefore be able to integrate into their hosts as prophages.

Most notably, two *Haemophilus*-targeting vOTUs carrying *neuA* were detected in 5 of 68 *Haemophilus*-positive samples (7.4%) (Figure 6A). Amongst *Haemophilus*-colonised patients, those harbouring either virulence-factor carrying phage showed dramatically elevated bacterial abundance compared to those without (median 37.3% vs 0.46%, 82-fold increase; mean 45.3% vs 3.7%, 12-fold increase; Wilcoxon test p = 5.5×10⁻⁴). This association was significant in COPD patients (4 of 46 colonised patients, p = 8.7×10⁻⁴), with only one instance detected in a healthy control (Figure 6B).

**Figure 6.**
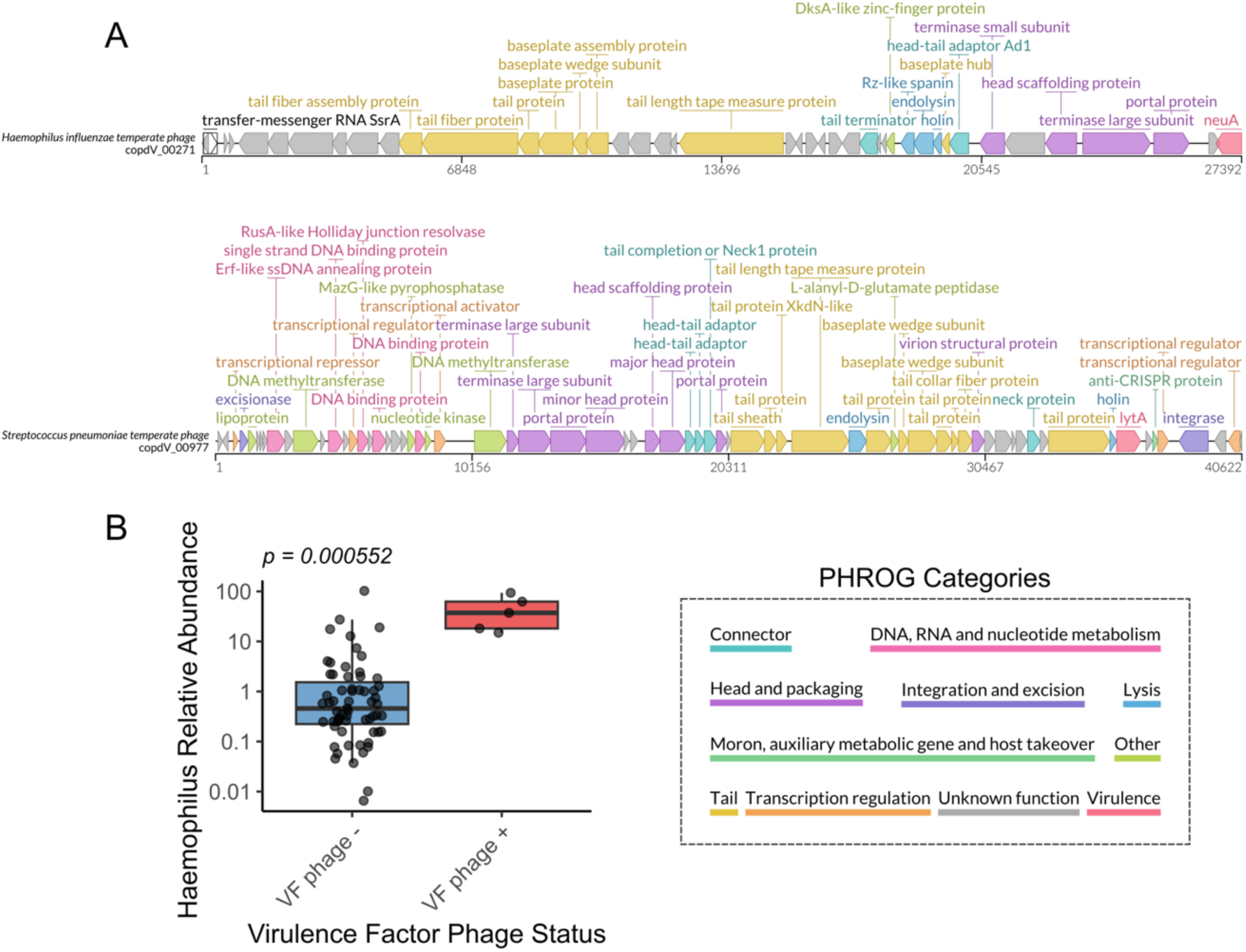
Virulence factor-encoding phages associate with elevated pathogen abundance. (**A**) Genome maps of two virulence factor-encoding vOTUs predicted to infect key COPD pathogens. Top: vOTU targeting *Haemophilus influenzae* carrying *neuA* (acylneuraminate cytidylyltransferase). Bottom: vOTU targeting *Streptococcus pneumoniae* carrying *lytA* (autolysin). Open reading frames (ORFs) are coloured by PHROG functional category. Scale bars indicate genome length in bases. (**B**) *Haemophilus* abundance in colonised patients with and without *neuA*-encoding phages. Box plots show median and interquartile range; individual samples shown as points coloured by disease status. Among 68 *Haemophilus*-positive samples, those harbouring virulence phages (n = 5) showed 82-fold higher median bacterial abundance compared to those without (n = 63). Statistical significance determined by Wilcoxon rank-sum test.

These virulence-factor-encoding vOTUs appeared across different clinical groups (three Group A, one Group B, one healthy control). The strong ecological coupling suggests either phage-mediated enhancement of bacterial fitness through virulence factor provision, or preferential phage replication in high-density bacterial populations. However, the low prevalence limits broader conclusions about their role in COPD pathogenesis.

## Discussion

This re-analysis of the Yan et al. COPD metagenome dataset^18^ reveals fundamental alterations in viral ecology that parallel and potentially contribute to disease progression. By applying specific virome-focused bioinformatic approaches to bulk metagenomes, we demonstrate that bacteriophage diversity loss represents a previously unrecognised feature of COPD correlating with disease severity and varying by inflammatory phenotype.

The progressive viral diversity decline from healthy controls through increasing COPD severity suggests viral community disruption may be integral to pathogenesis. This mirrors observations in inflammatory bowel disease and HIV infection, where reduced virome diversity correlates with disease activity^23-25^. The threshold effect—significant viral depletion manifesting primarily in GOLD stages 3-4—aligns with COPD as a disease of accumulated insults eventually overwhelming homeostatic mechanisms. Complete absence of detectable DNA viruses in some patients’ bulk metagenomes suggests profound ecological collapse, though this heterogeneity likely reflects varying disease trajectories, therapeutic histories, or sampling limitations.

Our finding that frequent exacerbators (Group E) uniquely exhibit decoupled viral-bacterial diversity relationships provides novel insight into this high-risk phenotype. Whilst viral and bacterial communities typically show coordinated responses, the absence of correlation in Group E suggests fundamentally altered microbial ecology. This may result from repeated antibiotic exposure during exacerbations, creating asynchronous selective pressures, or from the inflammatory milieu—characterised by neutrophil influx and oxidative stress—differentially affecting phage and bacterial survival. These findings have therapeutic implications: microbiome-targeted interventions in frequent exacerbators may need to consider viral communities independently from bacterial hosts.

The disproportionately lower abundance of phages targeting oral and anaerobic bacteria compared to their bacterial hosts adds complexity to understanding COPD dysbiosis. The 40-fold lower abundance of *Porphyromonas* phages despite only 4-fold lower bacterial abundance suggests factors beyond simple host availability. This could represent completed Kill-the-Winner cycles leaving resistant bacterial subpopulations, direct inflammatory effects on phage stability, or altered bacterial physiology reducing phage susceptibility. These differential patterns—lower abundance of phages targeting health-associated bacteria but unchanged pathogen-associated phages—could fundamentally alter community dynamics by removing phage-mediated control that normally limits pathogen expansion.

Persistent Kill-the-Winner dynamics across all COPD phenotypes, albeit with varying intensity, indicates fundamental ecological principles remain operative even in disrupted communities. The weakening of Kill-the-Winner dynamics in severe disease (GOLD 4) suggests deteriorating ecological regulation as communities simplify, with important implications for potential phage therapy approaches that must account for altered predator-prey dynamics and potentially reduced efficacy in advanced disease.

The acute viral events—both eukaryotic infections and bacteriophage blooms— observed in five patients warrant consideration. The phage blooms, characterised by single bacteriophages dominating the viral community (>80% relative abundance), likely represent prophage induction events or rapid lytic amplification during bacterial population expansions. These extreme viral loads coincided with markedly reduced diversity, suggesting periodic boom-bust cycles that may contribute to the overall viral community instability observed in COPD.

Detection of virulence factor-encoding phages, particularly the *neuA*-carrying *Haemophilus* phages associated with dramatically elevated bacterial abundance, provides direct evidence of phage-mediated virulence enhancement in COPD lungs. To our knowledge, this represents the first demonstration of virulence factor-encoding bacteriophages in the human respiratory tract, paralleling the paradigm established by *Vibrio cholerae*, where prophage-encoded cholera toxin transforms an otherwise harmless bacteria into a pathogen^12^.

Several limitations merit consideration. The cross-sectional design prevents causal inference about whether viral depletion drives or results from disease progression, and precludes assessment of how treatments—particularly antibiotics and corticosteroids— shape viral-bacterial dynamics over time. Analysis of bulk metagenomes rather than dedicated viromes likely underestimates viral diversity, particularly for low-abundance phages^26^. However, this approach captures both integrated prophages and free viruses, providing insight into temperate phage populations that may contribute to bacterial metabolism^27^. Our focus on DNA viruses excludes RNA viruses, established contributors to COPD exacerbations^7^. Additionally, whilst sputum sampling captures clinically relevant communities, it may not fully represent lower airway ecology. Small sample sizes for some inflammatory endotypes, particularly paucigranulocytic, limit generalisability for these groups.

Future research should establish temporal relationships between viral depletion and disease progression through longitudinal studies. Incorporating metatranscriptomics would reveal whether detected prophages are actively expressed, providing crucial insight into lysogenic-lytic switches and real-time phage-bacteria interactions that shape disease progression. Dedicated virome sequencing, including RNA viruses, would provide comprehensive viral characterisation. Investigation of common COPD therapeutics’ effects on viral communities could inform treatment strategies, whilst mechanistic studies examining how specific medications—particularly inhaled corticosteroids and antibiotics—shape viral ecology could guide targeted approaches. Exploring functional consequences of viral alterations, including effects on bacterial virulence, antibiotic resistance, and host immunity, could reveal novel therapeutic targets.

In conclusion, this analysis establishes alterations in bacteriophage ecology as a previously unrecognised feature of COPD pathobiology. Progressive viral diversity loss, disrupted viral-bacterial relationships in frequent exacerbators, and selective depletion of commensal-associated phages represent fundamental changes in lung microbial ecology likely contributing to disease progression. These findings highlight the importance of considering viral communities alongside bacteria in understanding respiratory dysbiosis and suggest comprehensive microbiome-targeted therapies should account for the viral component of the lung ecosystem.

## Methods

### Read QC and Assembly

Paired end fastq files were processed using fastp v0.24.0 with --detect_adapter_for_pe and -q 30^28^. Human reads had already been removed in the original analysis by mapping to human genome GRCh38 using BWA v.0.7.17^18,29^. Library sizes were calculated using the count command within SeqFu v1.22.3^30^. Samples were individually assembled using metaSPAdes v4.0.0 with default parameters^31^. Assembly graphs were used as input for Phables v1.4.1 using default parameters^32^, that is known to increase the recovery of complete viral genomes from human samples^33^.

### Viral Mining

Viral contigs were predicted using geNomad v1.8.1^34^. Predicted viruses were processed using CheckV v1.0.3 end_to_end ^35^. Putative viruses with complete ≥ 50% (equivalent to CheckV “Medium-quality) or length ≥ 10Kb were retained and de-replicated to form viral operational taxonomic units (vOTUs) following Minimum Information about an Uncultivated Virus Genome (MIUViG) standards (95% ANI over 85% length) using blast v2.14.0+ alongside the anicalc.py and aniclust.py scripts described in the CheckV documentation (https://bitbucket.org/berkeleylab/checkv/src/master/)^35-37^.

### vOTU Annotation

Functional annotation of vOTUs was performed using Pharokka v1.7.4 with -g prodigal-gv that uses tRNAscan-SE 2.0, Aragorn, CRT, PHROGs, VFDB, CARD, MMseqs2, PyHMMER, INPHARED and MASH^38-49^. Subsequent annotation was performed using PHOLD v0.2.0 (https://github.com/gbouras13/phold) that implements Foldseek and ProstT5^50,51^. Host prediction was performed using iPHoP v1.3.3^52-56^ and lifestyle was predicted using PhaTYP^57^. Family-level taxonomic assignments were extracted from geNomad and further taxonomic assignments were made using taxMyPhage v03.3^58^.

### Community Profiling

Reads were mapped to vOTUs using bowtie2 v2.5.4 with --non-deterministic and -- maxins 2000 with output piped to samtools v1.21 to produce a sorted bam file for each sample^59,60^. Abundance was calculated as CPM (analogous to the RNA-Seq measure of TPM) using CoverM contig v0.7.0 with --min-read-aligned-percent 90 and --min-covered-fraction 0.75^61^. Bacterial community composition was assessed using MetaPhlAn v4.1.1 with bowtie2 v2.5.4, -t rel_ab, -x mpa_vJun23_CHOCOPhlAnSGB_202307, -- ignore_eukaryotes, and --unclassified_estimation^59,62^. Diversity indices for bacteria were calculated using the genus level predictions and for viruses using CPM of vOTUs.

### Statistical Analysis

All statistical analyses were performed in R (v4.2.1) using the vegan package (v2.6-4) for ecological analyses. Alpha diversity metrics (Shannon diversity, Inverse Simpson, and observed richness) were calculated for both viral and bacterial communities. Due to non-normal data distributions and unequal sample sizes, non-parametric tests were employed throughout. Differences in diversity metrics across clinical groups (ABE classification, GOLD stages, and inflammatory endotypes) were assessed using Kruskal-Wallis tests, followed by Dunn’s post-hoc tests with Benjamini-Hochberg correction for multiple comparisons. Direct comparisons between two groups used Wilcoxon rank-sum tests. Correlations between continuous variables (viral diversity, bacterial diversity, eosinophil counts, viral load) were evaluated using Spearman’s rank correlation coefficient.

Beta diversity analysis was performed on viral abundance matrices using Bray-Curtis dissimilarity. Community structure differences between clinical groups were tested using permutational multivariate analysis of variance (PERMANOVA; adonis2 function, 999 permutations). To assess whether significant PERMANOVA results reflected true location differences or dispersion effects, we tested for homogeneity of multivariate dispersions using PERMDISP (betadisper and permutest functions, 999 permutations). Nested PERMANOVA was employed to first test overall COPD versus healthy differences, followed by within-COPD group comparisons. Pairwise PERMANOVA comparisons between groups were conducted with Benjamini-Hochberg correction for multiple testing.

To examine viral-bacterial community relationships, Mantel tests (999 permutations) were performed between Bray-Curtis dissimilarity matrices calculated from viral and bacterial abundance data. Partial Mantel tests controlled for potential confounders including disease status and GOLD stage. Procrustes analysis provided an independent assessment of community correlation through superimposition of ordinations. Environmental fitting (envfit function, 999 permutations) identified clinical variables significantly associated with viral community structure. Statistical significance was defined as p < 0.05, with marginally significant results (0.05 < p < 0.10) noted where relevant.

## Data Visualisation

Figures were produced using ggplot2 in R^63,64^. Genome maps were produced using LoVis4u^65^. Synteny comparisons were produced using Clinker^66^.

## Data Availability

No primary data was generated as part of this study. The vOTU catalogue and supplementary tables have been made available on Figshare at: https://figshare.com/s/eb8f56f9cc1e0a2cbcdc.

## Acknowledgments

We gratefully acknowledge all authors of the original publication that produced this dataset. We also acknowledge the contributions of all patients and participants that contributed their samples, and the clinicians that are overseeing treatment of these patients. This work was supported by the NIHR Leicester Biomedical Research Centre and the NIHR Leicester Clinical Research Facility. The views expressed are those of the author(s) and not necessarily those of the NHS, the NIHR or the Department of Health and Social Care.

R.A.C. and E.M.A. are funded through the Biotechnology and Biological Sciences Research Council (BBSRC) grant Bacteriophages in Gut Health BB/W015706/1. E.M.A. gratefully acknowledges the support of the BBSRC; this research was funded by the BBSRC Institute Strategic Programme Food Microbiome and Health BB/X011054/1 and its constituent projects BBS/E/F/000PR13631 and BBS/E/F/000PR13633; and by the BBSRC Institute Strategic Programme Microbes and Food Safety BB/X011011/1 and its constituent projects BBS/E/F/000PR13634, BBS/E/F/000PR13635 and BBS/E/F/000PR13636. A.M. was funded by MRC-CLIMB (MR/L015080/1) and CLIMB-BIG-DATA (MR/T030062/1), with bioinformatics analysis carried out on infrastructure provided by MRC-CLIMB and CLIMB-BIG-DATA. Y.Y., Z.L., F.W., R.C., and Z.W. would like to gratefully acknowledge National Key R&D Program of China (2022YFA1304300 to Z.W.), National Natural Science Foundation of China (82495200, 82495201, 32170109 to Z.W.), Major Project of Guangzhou National Laboratory (GZNL2023A02001 to Z.W.), the Noncommunicable Chronic Diseases-National Science and Technology Major Project (2024ZD0528400 to Z.W.)

## Contributions

R.A.C. and C.E.B. conceived the idea. Y.Y., Z.L., F.W., R.C. and Z.W. provided data and materials. R.A.C., A.J.P., A.T., M.R.J.C, A.D.M, and C.E.B. analysed the data. R.A.C. drafted the manuscript. R.A.C., A.J.P., A.T., Z.W., E.M.A., M.R.J.C, A.D.M, and C.E.B. edited the manuscript. All authors read and approved the final manuscript.

## Ethics Declarations

The authors have no competing interests to declare. No additional ethics were required for this re-analysis of previously published data.

## Supplementary Materials

An excel workbook is available with four supplementary tables. **Supplementary Table 1:** Sample data. **Supplementary Table 2:** vOTU characteristics. **Supplementary Table 3:** vOTU abundance table (CPM). **Supplementary Table 4:** Bacterial abundance table.

